# GLI1 facilitates rheumatoid arthritis by collaborative regulation of DNA methyltransferases

**DOI:** 10.1101/2023.02.07.527456

**Authors:** Gaoran Ge, Qianping Guo, Ying Zhou, Wenming Li, Wei Zhang, Jiaxiang Bai, Qing Wang, Huaqiang Tao, Wei Wang, Zhen Wang, Minfeng Gan, Yaozeng Xu, Huilin Yang, Bin Li, Dechun Geng

## Abstract

Rheumatoid arthritis (RA) is characterized by joint synovitis and bone destruction, the etiology of which remains to be explored. Overactivation of M1 macrophages and osteoclasts has been thought a direct cause of joint inflammation and bone destruction. Glioma-associated oncogene homolog 1 (GLI1) has been revealed to be closely linked to bone metabolism. In this study, GLI1-expression in synovial tissue of RA patients showed to be positively correlated with RA-related scores and was highly expressed in collagen-induced arthritis (CIA) mouse articular macrophage-like cells. The decreased expression and inhibition of nuclear transfer of GLI1 downregulated macrophage M1 polarization and osteoclast activation, the effect of which was achieved by modulation of DNA methyltransferases (DNMTs) via transcriptional regulation and protein interaction ways. By pharmacological inhibition of GLI1, the proportion of proinflammatory macrophages and the number of osteoclasts were significantly reduced, and the joint inflammatory response and bone destruction in CIA mice were alleviated. This study clarified the mechanism of GLI1 in macrophage phenotypic changes and activation of osteoclasts, suggesting potential applications of GLI1 inhibitor in the clinical treatment of RA.

## 1. Introduction

Rheumatoid arthritis (RA) is a common chronic inflammatory disease that currently affects approximately 75 million people worldwide. In addition to arthritis symptoms, RA can eventually lead to disability or even death [1, 2]. Although the mechanism of RA is not clear, it is generally believed that synovial inflammation and bone erosion are the direct factors causing joint damage. Therefore, unravelling the molecular pathways that underlie immune regulation and bone destruction are of major interest to better understand the pathophysiology of RA and to design new approaches to achieve a therapy for this severe joint disease.

Persistent activation of immune cells leads to the progression of symptoms such as synovitis in RA. This immune-mediated disorder involves, both innate and adaptive cellular compartments as well as their dysregulated cytokine production. In addition, the pathological process of RA is promoted through the synergistic action of the cellular resident in the bone and in joint compartments, such as osteoclast, chondrocyte and stromal cells are involved [3]. Within the innate immunity function that causes inflammation, macrophages have a key role. As a matter of fact, they contribute to the normal tissue homeostasis, working as antigen-presenting cells (APCs) to activate adaptive immunity, pathogen expulsion, resolution of inflammation and wound healing [4]. Classically activated macrophages (M1), which has been proven to be dominant in RA joints, causes joint erosion, secreting principally proinflammatory cytokines such as tumor necrosis factor α (TNF-α), interleukin 1β (IL-1β) and IL6, whereas alternatively activate macrophages (M2) contributes to tissue remodeling and repair via a high production of anti-inflammatory cytokines (mainly IL-10 and TGF-β) [4-6]. Another hallmark performs as irreversible bone destruction, which is leaded by osteoclasts. The RA microenvironment and those proinflammatory cytokines stimulate osteoclast formation, which has long been considered the only cell capable of absorbing bone matrix, and the subsequent degradation of bone and cartilage [7-9]. Consequently, effectively hindering the overactivation of osteoclasts is one of the keys to alleviate excessive bone resorption.

Hedgehog pathway, which has been shown to be closely involved in osteogenesis [10], is highly conserved and is critical for normal embryogenesis. The hedgehog protein family consists of hedgehog ligands binding the cell surface transmembrane receptor patch (PTCH) and functioned through posttranslational processing of glioma-associated oncogene homolog (GLI) zinc finger transcription factors [11, 12]. To date, three mammalian GLI proteins have been identified, among which GLI1 usually acts as a transcriptional activator. Many of the physiopathological processes involved with GLIs are complex and worth discussing. Relevant studies have shown that GLI1-activated transcription promotes the development of inflammatory diseases such as gastritis, and antagonizing GLI1 transcription can alleviate the inflammatory degradation of articular cartilage [13, 14]. Accordingly, these clues suggest that GLI1 may be involved in inflammation and bone erosion processes, driving us to explore its role and regulatory mechanisms in RA.

In this study, we investigated the association of GLI1 with RA as well as the critical role of GLI1 on pathological process of joint inflammation and bone destruction. It was confirmed that inhibition of GLI1 could reduce the inflammatory bone destruction in collagen-induced arthritis (CIA) mice. Mechanistically, the intranuclear translocation of GLI1 was shown to promote M1 polarization of macrophages and over activation of osteoclasts by collaborative regulation of DNA methyltransferases (DNMTs). In general, this study not only clarified the pivotal role and mechanism of GLI1 in joint inflammation and bone destruction of RA, but also suggested potential applications of GLI1’s inhibitor in the clinical treatment.

## 2. Results

### 2.1. GLI1 expression is elevated in human RA synovium, and selective inhibitor of GLI1 can alleviate joint bone destruction in CIA mice

To determine the relationship between GLI1 expression and the RA process, we collected synovial tissue from RA patients and non-inflammatory joint diseases donors and obtained informed consent from the patients. Compared with normal synovial tissue, synovium of RA patients with have a large number of inflammatory cell infiltration (**Fig. 1a**). Immunohistochemical (IHC) staining showed that there were more GLI1-positive cells in RA synovium than those in normal synovium (**Fig. 1b, c**). The results of western blotting further verified the upregulation of GLI1 in the joint synovium of RA patients (**Fig. S1a**). We performed a DAS28 score on the included patients to initially assess RA activity and then matched the relative expression of GLI1 with them, indicating that there was a linear relationship between GLI1 expression and RA activity to a certain extent (**Fig. S1b, c**). These results were of concern to us and implied that GLI1 plays a catalytic role in the pathological process of RA.

**Figure 1.**
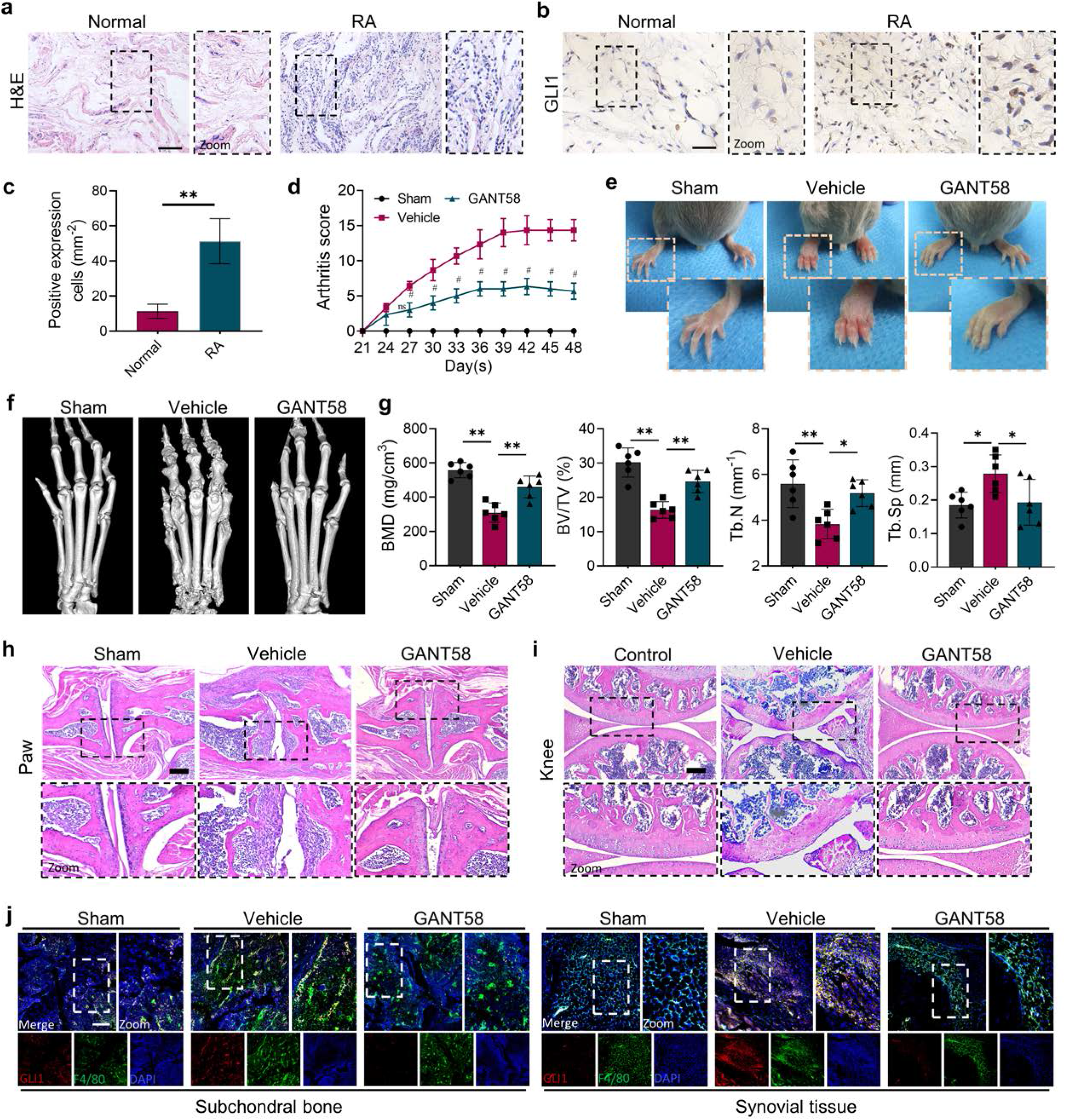
GLI1 is highly expressed in human RA synovium while regulating the pathological process of joint inflammatory bone destruction in CIA mice. **a** H&E staining of normal and RA human synovium. Scale bars = 200 μm. **b** IHC staining of GLI1 in normal and RA human synovium and **(c)** quantification of positive stained cells. Scale bars = 50 μm. n = 3. **d** Arthritis score of mice limbs. **e** Photos of mice paws. **f** Micro-CT scanning and 3D reconstruction of mouse paws. **g** Bone parameters of BV/TV, BMD, Tb.N, Tb.Th. **h, i** Images of H&E staining of mouse paw and knee joints. Scale bars = 200 μm. **j** Immunofluorescence co-staining image of F4/80 with GLI1 (Red: GLI1, green: F4/80, blue: DAPI). Scale bars = 200 μm. The in vivo results are presented as the mean ± SD of 6 mice per group. Data shown represent the mean ± SD. Statistical analysis was performed using one-way ANOVA test. *p < 0.05, **p < 0.01.

To verify this speculation, we constructed a collagen-induced arthritis (CIA) model using DBA mice and performed a therapeutic intervention with the GLI1 selective inhibitor GANT58. H&E staining results showed no hepatorenal toxicity of GANT58 (**Fig. S2**). After secondary immunization, CIA mice’s paws swelled rapidly and there was weight loss simultaneously, which could be reversed by GANT58 treatment (**Fig. S3a, b**). Based on the arthritis score, it could be observed that after 21 days of modeling, the scores of the mice in the model group increased significantly. Obviously, the arthritis scores of GANT58-treated mice were remarkably lower than those of mice in the vehicle group (**Fig. 1d**). We photographed the hind paws of mice in each group and found that CIA mice had obvious signs of arthritis, while mice in the GANT58 treatment group showed a slighter degree of swelling, which was similar to those of normal mice (**Fig. 1e**). Micro-CT was performed to scan the mouse paws and ankle joints. As shown in **Fig. 1f**, compared with sham group mice, vehicle group mice had more obvious bone destruction, and GANT58 treatment alleviated this effect. Through the analysis of bone parameters in the joint part of the mouse foot claw, we also found that the bone parameters were markedly better in the GANT58-treated mice than in the vehicle control mice (**Fig. 1g**). Based on these results, we performed histological sectioning and staining of mouse paw articular tissue as well as knee tissue to evaluate pathological histological changes in the joints. The results showed that the GANT58 intervention significantly reduced the degree of inflammatory cell infiltration and joint bone destruction (**Fig. 1h, i**; **Fig. S3c, d**). These evidences suggested that inhibiting the activity of GLI1 significantly reduced the occurrence and development of arthritis in CIA mice. However, it was not clear how the specific regulatory effect of GLI1 is. Thus, we performed immunofluorescence co-localization staining of the macrophage marker F4/80 and GLI1 both in Subchondral bone marrow cavity and synovial tissue and observed that the high expression of GLI1 in CIA mice was mostly located in macrophage-like cells (**Fig. 1j**). Therefore, our follow-up research mainly focused on the mechanism of GLI1 on the regulation of macrophage fate.

### 2.2. GLI1 affects the release of inflammatory cytokines by regulating the macrophage phenotype

To investigate the relationship between GLI1 expression and M1 macrophages in vitro, we first measured GLI1 protein expression by western blotting during M1 macrophage induction, which increased at 24 h after LPS and IFN-γ induction (**Fig. S4a**). Typically, when the GLI1-related pathway is activated, GLI1 is transported from the cytoplasm to the nucleus, which in turn exerts its transcriptional regulatory role. Therefore, we separately examined the amount of protein for GLI1 inside and outside the nucleus. As shown in **Fig. 2a** and **b**, after the LPS/IFN-γ intervention, although the amount within the cytoplasm has not changed significantly, the expression of GLI1 in nucleus increased to certain degrees, suggesting that GLI1 was activated and incorporated into the nucleus during the M1 polarization process to play a related regulatory role.

**Figure 2.**
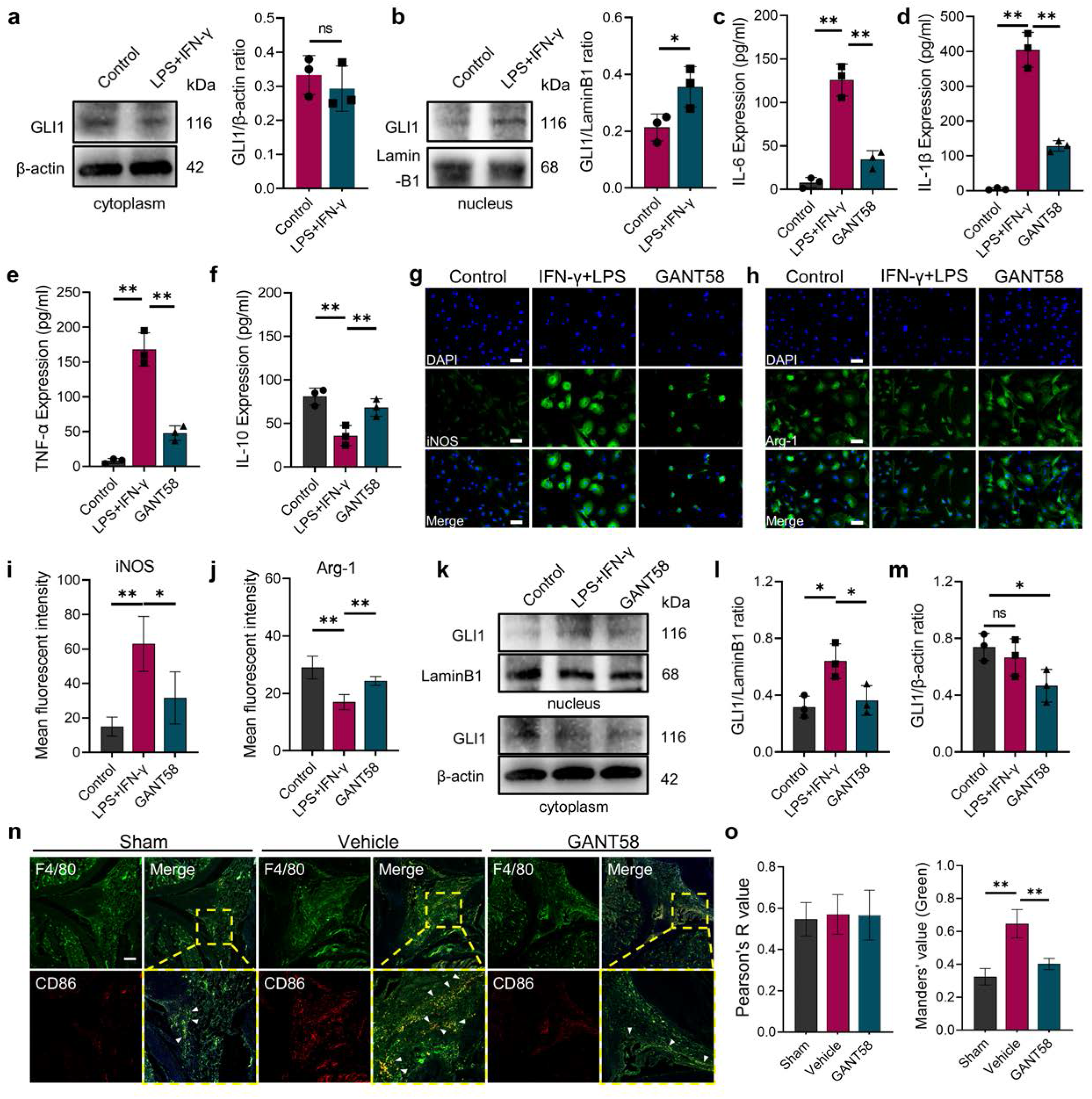
GLI1 plays an active role in M1 macrophage formation and the release of pro-inflammatory cytokines. **a, b** RAW264.7 cells were stimulated with LPS/IFN-γ for 24 h. The proteins in the cytoplasm and nucleus are isolated and extracted. GLI1 protein in the cytoplasm and nucleus was detected by western blot and grayscale value ratio to β-actin of western blot results was analyzed. n = 3. Statistical analysis was performed using Student’s t test. **c-f** Macrophages were stimulated with LPS/IFN-γ for 36 h. The supernatant was collected, and the volumes of IL-1β, IL-6, TNF-α and IL-10 were detected by ELISA. **g, h** Immunofluorescence staining of iNOS and arginase (Arg-1) during LPS/IFN-γ-induction. Scale bars = 25 μm. **i, j** Mean fluorescence intensity of immunofluorescence was analyzed using ImageJ. **k-m** RAW264.7 cells were stimulated with LPS/IFN-γ for 24 h, with or without GANT58 pretreatment. GLI1 protein in the cytoplasm and nucleus was detected by western blot and grayscale value ratio to β-actin of western blot results was analyzed. **n** Immunofluorescence staining of mouse joint tissue (green: F4/80, red: CD86). Scale bars = 200 μm. **o** Pearson colocation coefficient and CD86 positive quantitative analysis were performed with ImageJ. Statistical analysis was performed using one-way ANOVA test. Data shown represent the mean ± SD. *p < 0.05, **p < 0.01.

Given to this finding, we further utilize GANT58 to intervene in the macrophage polarization process. Cell viability was measured by CCK-8 assay before the in vitro using of GANT58, and the results showed that GANT58 had no significant inhibitory effect neither on BMMs and RAW264.7 cells until the concentration reached 40 μM (**Fig. S4b-e**). We treated cells with a concentration of 10 μM and found to reduce the expression of GLI1 (**Fig. S5a, b**). Subsequently, we performed a GANT58 intervention on M1-induced macrophages, collecting and detecting changes in inflammatory cytokine content in the cell supernatant. Stimulated by LPS and IFN-γ, macrophages release a large number of pro-inflammatory cytokines including IL-1β, IL-6, and TNF-α. After the inhibitory intervention of GLI1, the content of these cytokines was significantly reduced, while the release inhibition of IL-10 was alleviated to some extent (**Fig. 2c-f**). Immunofluorescence staining suggested that the GANT58 group exhibited fewer iNOS-positive cells and near-basic level of Arg-1 expression compared to cells in the LPS and IFN-γ treated groups (**Fig. 2g-j**), and the western blot results supported the same conclusion (**Fig. S6a, b**). In addition, we examined the expression and distribution of GLI1 in cells under different interventions and found that GANT58 significantly reduced the nucleation transport of GLI1 while inhibiting the pro-inflammatory state of cells (**Fig. 2k-m**). Subsequently, we performed immunofluorescence co-staining of F4/80 and CD86 in animal model tissues and compared with more CD86-positive M1 macrophages in the synovial tissue of mice in the CIA model group, the number of positive macrophages of the pro-inflammatory phenotype after GANT58 treatment was significantly reduced (**Fig. 2n, o**). These results confirmed that GANT58 was capable of reducing the maturation of M1 macrophages and the release of pro-inflammatory cytokines by affecting the activation of GLI1 both in vitro and in vivo.

### 2.3. The expression and intranuclear transport of GLI1 is involved in osteoclast activation

The over activation of osteoclast is the direct cause of bone destruction in RA. In previous studies, we have found that GLI1 is highly expressed in macrophage-like cells in the subchondral bone of the joints, which raised our concerns about GLI1 and osteoclasts. To investigate the role of GLI1 in osteoclast formation, we measured GLI1 protein expression of RAW264.7 3 days after RANKL induction (**Fig. 3a**). The results showed that after RANKL stimulation, more GLI1 was activated and transferred from the cytoplasm to the nucleus (**Fig. 3b, c**). Thus, in order to confirm the regulatory function of GLI1 in the differentiation of osteoclasts, we used GANT58 to intervene GLI1 activity during the RANKL induction. Original BMMs were isolated and cultured with M-CSF, followed by stimulating with RANKL, with or without GANT58 pretreatment. As a result, there was less osteoclast formation in the GANT58-treated group than that in the RANKL stimulation group (**Fig. 3d**). Osteoclast formation is accompanied by high expression of NFATc1. It could be observed by immunofluorescence staining that the number of NFATc1-positive cells decreased significantly after GANT58 treatment (**Fig. 3e**). We further performed the same treatment on macrophage cell line RAW264.7 cells. Tartrate-resistant acid phosphatase (TRAP) and F-actin staining confirmed that the intervention of GANT58 greatly reduced the formation of mature osteoclasts as well (**Fig. 3f, g**). In addition, we extracted the total protein from RAW264.7 cells with different treatment and performed western blot experiments. During osteoclast induction, GANT58 significantly inhibited the expression of markers related to osteoclast function, including NFATc1, CTSK and MMP9 (**Fig. 3h**). In this process, we also found that GANT58 hindered the activation of GLI1 translocation into the nucleus (**Fig. 3i**). To further verify the effect of GLI1 osteoclast activation in RA, we performed TRAP staining of subchondral bone in mice of different groups. The results showed that there were more osteoclast formations around the trabeculus in the CIA model group. After GANT58 treatment, the number of osteoclasts decreased, and the trabecular morphology was close to the normal tissue level (**Fig. 3j**; **Fig. S6c**). In summary, these data suggested that the activity of GLI1 may facilitate osteoclast formation.

**Figure 3.**
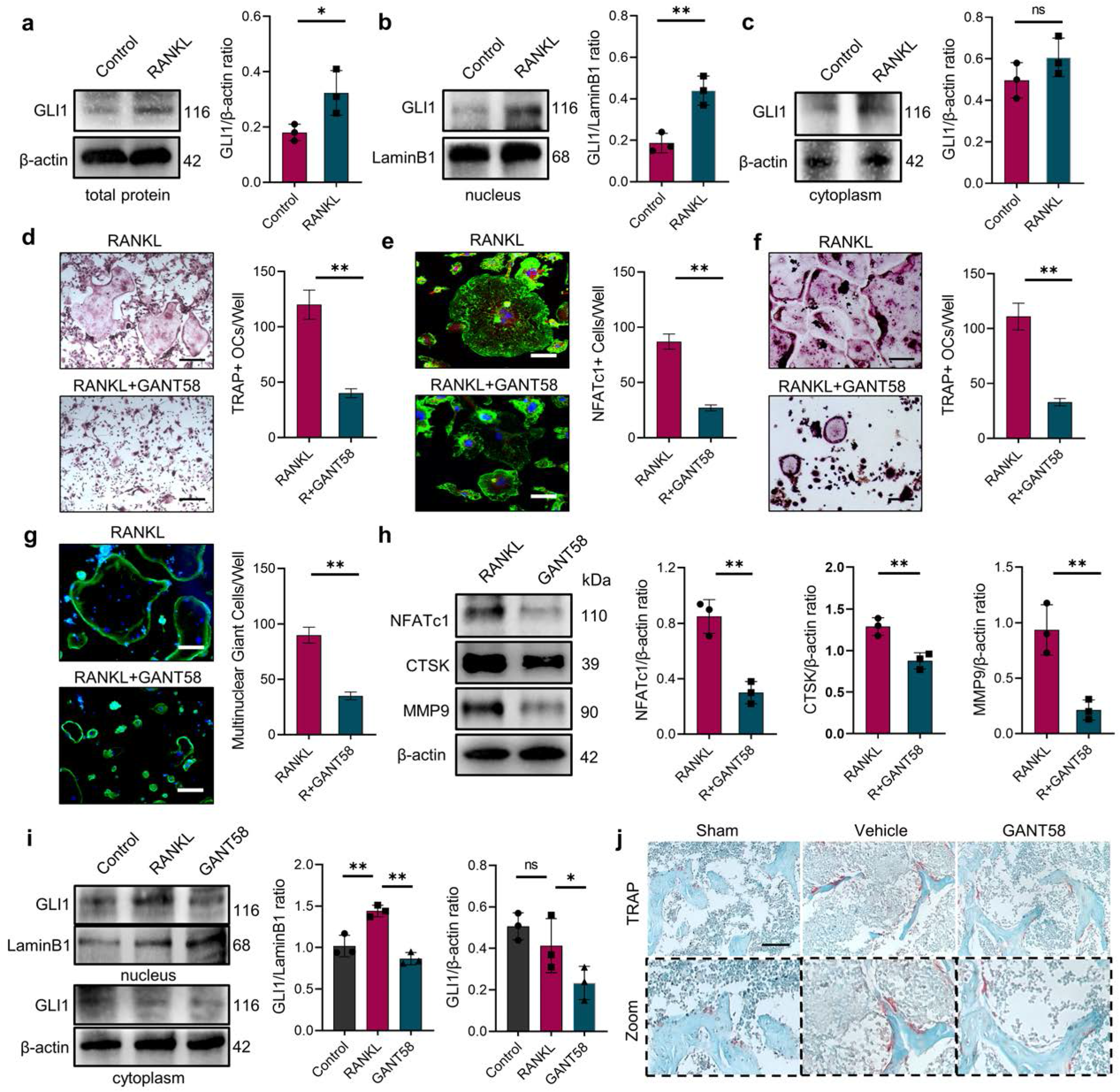
GLI1 regulates the formation of osteoclasts in vivo and in vitro. **a-c** RAW264.7 cells were stimulated by RANKL. Total GLI1 protein and GLI1 in the cytoplasm and nucleus was detected by western blot during the induction of osteoclasts and the grayscale value ratio to β-actin of western blot results was analyzed. n = 3. **d** TRAP staining of BMMs with RANKL induction and TRAP positive osteoclasts quantity per well. Scale bars = 25 μm. **e** F-actin and NFATc1 immunofluorescence staining of RANKL-stimulated BMM-derived osteoclasts and NFATc1-positive cells quantity per well. Scale bars = 25 μm. **f** TRAP staining of RAW264.7 with RANKL induction and TRAP positive osteoclasts quantity per well. Scale bars = 25 μm. **g** F-actin immunofluorescence staining of RANKL-stimulated RAW264.7-derived osteoclasts and multinucleated giant cells quantity per well. Scale bars = 25 μm. **h** RAW264.7 cells were stimulated by RANKL for 3 days. Western blot analysis of NFATc1, CTSK, and MMP9 was performed and grayscale value ratio to β-actin of western blot results. n = 3. Statistical analysis was performed using Student’s t test. **i** RAW264.7 cells were stimulated with RANKL for 3 days, with or without GANT58 pretreatment. GLI1 protein in the cytoplasm and nucleus was detected by western blot and grayscale value ratio to β-actin of western blot results was analyzed. Statistical analysis was performed using one-way ANOVA test. **j** TRAP staining of mouse joints subchondral bone tissue. Scale bars = 100 μm. Data shown represent the mean ± SD. *p < 0.05, **p < 0.01, ns= no significance.

### 2.4. GLI1 regulates the expression of DNMTs in distinct ways during the different fates of macrophages

As a nuclear transcription factor, GLI1 exerts an active effect through nuclear entry. In order to explore the potential downstream regulation mechanism of GLI1, RNA sequencing (RNA-seq) on the macrophages before and after GLI1 intervention was performed then to observe gene expression changes. The seq data showed that more genes were down-regulated than up-regulated in GANT58 treated cells (**Fig. S7a**). Among these differentially altered genes, we revealed through Gene Ontology (GO) analysis that GANT58’s intervention in GLI1 affected multiple biological processes including macrophage chemotaxis and macrophage cytokine production (**Fig. 4a**). The results of the Kyoto Encyclopedia of Genes and Genomes (KEGG) enrichment analysis showed that these down-regulated genes were involved in the development of human diseases such as rheumatoid arthritis, as well as organismal systems such as osteoclast differentiation (**Fig. 4b**). These evidences confirmed our previous results. Specifically, GANT58 reduced some of the osteoclast and inflammation-related genes in the cell resting state.

**Figure 4.**
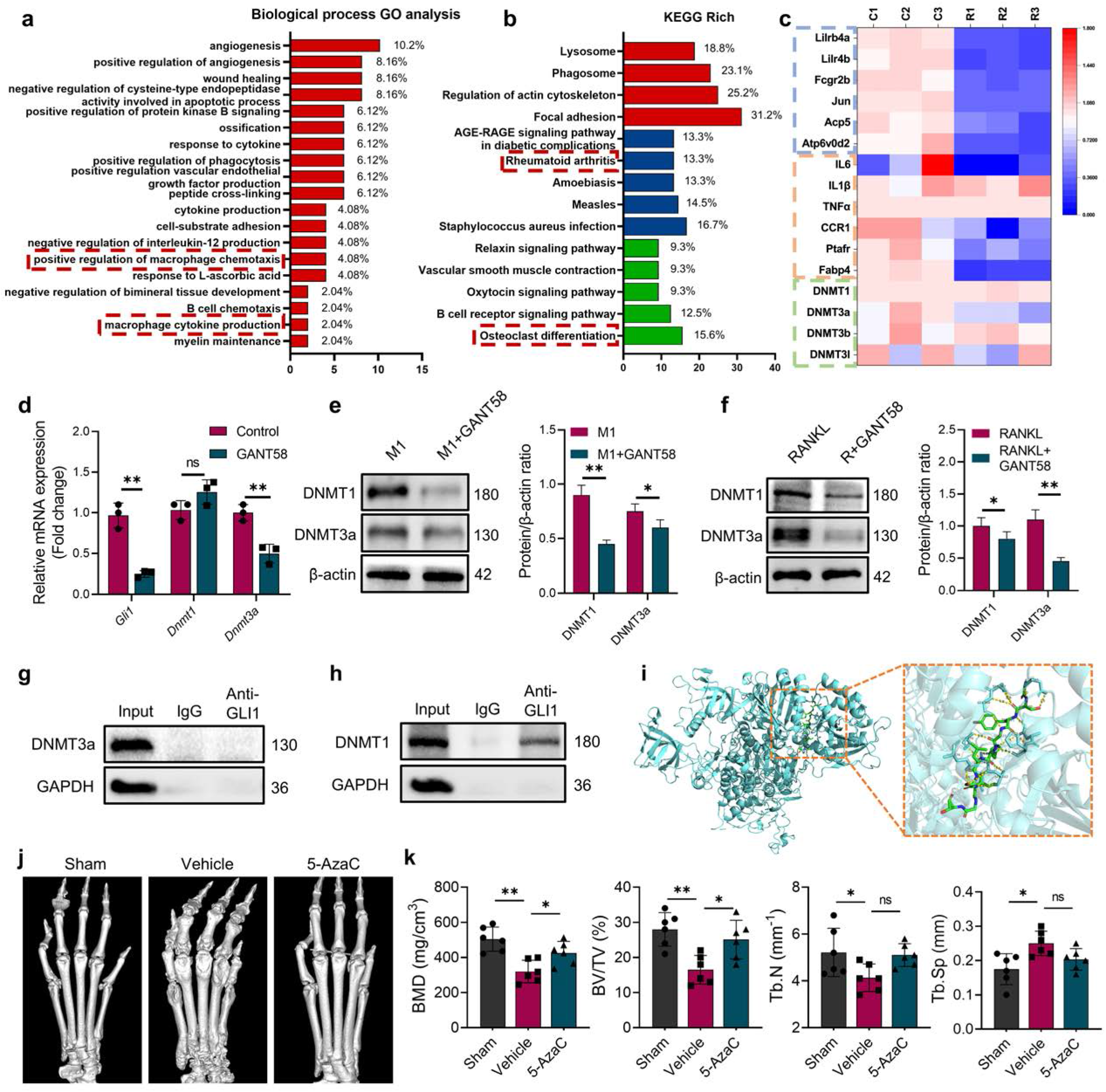
DNA methyltransferases might be a regulatory target downstream of GLI1. **a** Biological process GO analysis of RNA-seq results for macrophages with or without GANT58 treatment. **b** KEGG rich analysis of RNA-seq results. **c** Heat map of parts of the relevant gene transcriptional expressions (C = control; R = GANT58; red: increased expression; blue: decreased expression). **d** Relative mRNA expression of Gli1, Dnmt1 and Dnmt3a in macrophages with or without GANT58 treatment. Statistical analysis was performed using two-way ANOVA test. **e** RAW264.7 cells were stimulated by LPS and IFN-γ for 24 h, with or without GANT58 co-intervention. Western blot results of DNMT1 and DNMT3a protein expression and grayscale value ratio to β-actin of western blot results. n = 3. **f** RAW264.7 cells were stimulated by RANKL for 3 days, with or without GANT58 co-intervention. Western blot results of DNMT1 and DNMT3a protein expression and grayscale value ratio to β-actin of western blot results. n = 3. Statistical analysis was performed using two-way ANOVA test. **g, h** Co-IP detection of protein binding between GLI1 and DNMT1/DNMT3a. n = 3. **i** Protein–protein interface interaction of GLI1 and DNMT1 with PyMOL. **j** Micro-CT scanning and 3D reconstruction of mouse paws. **k** Bone parameters of BV/TV, BMD, Tb.N, Tb.Th. n = 6. Statistical analysis was performed using one-way ANOVA test. Data shown represent the mean ± SD. *p < 0.05, **p < 0.01, ns= no significance.

It is worth noting that among these genes, the expression of *Dnmt3a* of the DNA methyltransferase family also showed a decreasing trend (**Fig. 4c**). DNA methylation is an important epigenetic modification way to regulate gene expression, which is activated by DNMTs [15]. In mammals, DNMT1 and DNMT3a are the main methylation regulatory enzymes responsible for the de novo methylation and methylation maintenance of DNA. Furthermore, the results of bioinformatics analysis in GeneMANIA showed a latent relationship between GLI1 and DNMTs (**Fig. S7b**). Through protein detection of human synovial tissues, we found that DNMT1 and DNMT3a were more expressed in RA patients (**Fig. S8**). Thus, through further RT-qPCR validation, we found that *Dnmt3a* had a clear inhibition after the GANT58 intervention, but the expression of *Dnmt1* did not appear to change significantly (**Fig. 4d**). To this end, we wondered whether there will be another outcome in different induction states. Therefore, we further examined DNMT1 and DNMT3a expression changes during M1 macrophages and osteoclast induction processes with GANT58 intervention or not. It is interesting to note that although GANT58 only affected the expression of the DNMT3a gene in the resting state, the western blot results suggested that GANT58 reduced the expression of DNMT1 during M1 macrophage induction and the expression of DNMT3a during osteoclast induction, respectively (**Fig. 4e, f**). The expression and localization of the proteins detected by immunofluorescence staining demonstrated that DNMT1 and DNMT3a were highly expressed in the nucleus under LPS/IFN-γ and RANKL stimulation, and GANT58 reduced their nuclear expression in both RAW264.7 cells and BMMs (**Fig. S9a, b**). Co-ip protein binding experiments have found that it was DNMT1, but not DNMT3a protein could be pulled down by the antibody of GLI1, suggesting that there was a direct binding between GLI1 and DNMT1 (**Fig. 4g, h**). Accordingly, we used ZDOCK server to find possible binding sites and performed the docking of protein–protein interface interaction of GLI1 and DNMT1 with PyMOL (**Fig. 4i**; **Table S2**). Overall, the above validation results confirmed that GLI1 might have a regulatory effect on DNMTs during different induction processes of macrophages, and this regulation mode were not exactly the same.

Based on the above results, we treated CIA mice with 5-azacytidine (5-AzaC), an inhibitor of DNMTs, as a therapeutic intervention to observe the potential regulatory effect of DNMTs on RA. As shown in **Fig. 4j** and **k**, 5-AzaC significantly alleviated joint bone destruction in CIA mice and improved bone parameter results. This suggested that DNMTs might be involved in the RA development process, further suggesting its potential regulatory relationship with GLI1.

### 2.5. GLI1 regulates the proinflammatory phenotype of macrophage by affecting the expression of DNMT1

As an important member of the DNMT family, apart from being involved in the occurrence of tumor diseases, DNMT1 has also been shown to be associated with certain inflammatory diseases [16]. By staining mouse joints for IHC, we could find that DNMT1 was more expressed in the CIA model group, while its expression was reduced in the GANT58 treatment group (**Fig. 5a, b**; **Fig. S10a, b**). We first observed the alteration of DNMTs during in vitro macrophage polarization induction. The western blot results showed that DNMT1 was highly expressed during M1 macrophage induction than in M2 induction (**Fig. 5c-e**). To investigate the effect of DNMTs on macrophages, the DNMT3a-specific inhibitor theaflavin-3,3’-digallate (TF3) and the DNMT1-specific inhibitor procainamide (PR), were utilized for DNMTs intervention. The CCK-8 results showed that TF3 did not inhibit cell proliferation at concentrations ranging from 0 to 5 μM in either BMMs or RAW264.7 cells and no obvious proliferative toxic effect of PR could be seen on BMMs or RAW264.7 cells (**Fig. S11a-d**). We used different interventions to treat LPS/IFN-γ-induced macrophages and performed immunofluorescence staining to observe the expression of iNOS in proinflammatory M1 macrophages. An increase in the number of iNOS positively stained cells was observed after stimulation with LPS and IFN-γ, and similar to the effect of GANT58, the inhibition of DNMT1, but not DNMT3a was able to reduce M1 macrophage numbers (**Fig. 5g**). According to this result, we overexpressed *Dnmt1* and *Dnmt3a* by lentiviral transfection (GenePharma, Suzhou, China) (**Fig. 5f, g**). The results showed that the inhibitory effect of GANT58 on M1 macrophages was reversed by *Dnmt1* overexpression (OE) (**Fig. 5h-j**). In addition, we measured the mRNA expression of related cytokines, including the proinflammatory cytokines IL-1β, IL-6, and TNF-α and the anti-inflammatory cytokine IL-10, and found the same trend as that of the fluorescent staining results (**Fig. S12a-d**). These findings suggest that the regulatory effects of GLI1 on proinflammatory cells are mediated by DNMT1 (**Fig. 5k**).

**Figure 5.**
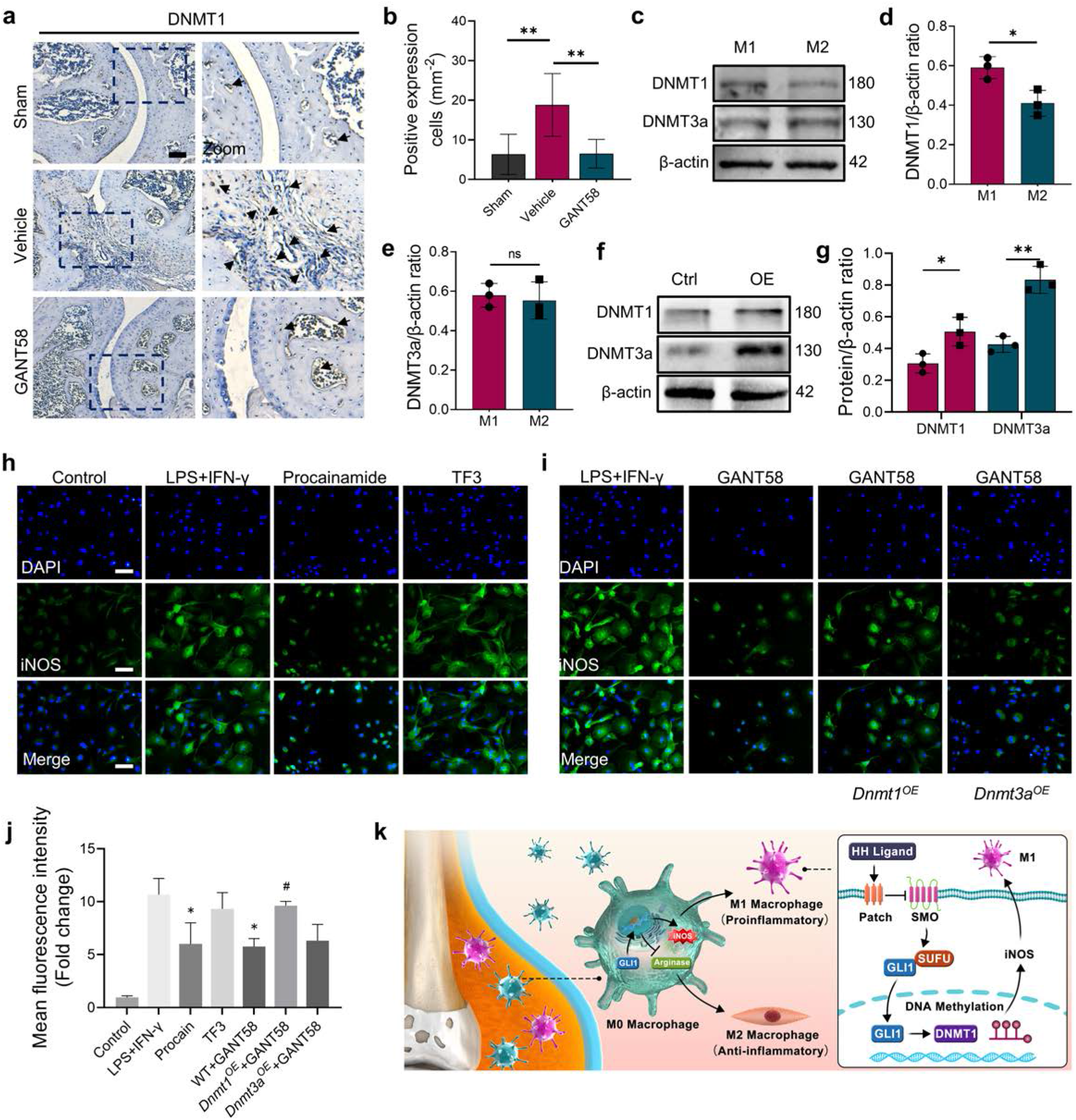
GLI1 regulated the expression of DNMT1, which affects the macrophage phenotypes and release of inflammatory cytokines. **a** IHC staining of DNMT1 and **b** quantification of positive cell numbers in mouse ankle joints. n = 5. Scale bars = 100 μm. Statistical analysis was performed using one-way ANOVA test. **c** RAW264.7 cells were stimulated by LPS/IFN-γ and IL-4. Western blot detection of DNMTs protein expression at 24 h. **d** DNMT1 grayscale value ratio to β-actin of western blot results. n = 3. *p < 0.05. **e** DNMT3a grayscale value ratio to β-actin of western blot results. n = 3. ns = no significance. **f** RAW264.7 cells were transfected with Dnmt3a and Dnmt1 overexpression lentiviruses. Western blot analysis of DNMT1 and DNMT3a protein expression. **g** Greyscale value ratio to β-actin of western blot results. n = 3. Statistical analysis was performed using Student’s t test. *p < 0.05, **p < 0.01. **h, i** BMMs were cultured and stimulated with LPS/IFN-γ in the presence of different interventions. Immunofluorescence staining of iNOS in LPS/IFN-γ-induced BMMs and **(j)** relative quantitative analysis of mean fluorescence intensity. Scale bars = 10 μm. n = 3. **k** Schematic diagram of the regulation mechanism. Statistical analysis was performed using one-way ANOVA test. Data shown represent the mean ± SD. *p < 0.05 compared with LPS/IFN-γ group; #p < 0.05 compared with WT+GANT58 group.

### 2.6. Overexpression of DNMT3a reverses osteoclast-forming disorders under GANT58 intervention

IHC staining of tissue specimens showed that similar to the GLI1 expression trend, DNMT3a expression was elevated in the CIA model group, which could be decreased by GANT58 (**Fig. S13a-c**). Meanwhile, western blot results showed that DNMT3a was also highly expressed during RANKL induction, while DNMT1 had no obvious change (**Fig. 6a, b**). To investigate the effect of DNMT3a and DNMT1 on osteoclast formation, the DNMT3a-specific inhibitor TF3 and the DNMT1-specific inhibitor PR were used for intervention in osteoclast induction. We first treated both BMMs and RAW264.7 cells with TF3 and PR simultaneously during osteoclast induction. As a result, RANKL-induced osteoclast formation was strongly inhibited by TF3 but not PR treatment (**Fig. 6d-g**). To verify whether the regulatory effect of GLI1 on osteoclasts was related to DNMTs, we pretreated wild-type (WT) RAW264.7 cells, *Dnmt3a* and *Dnmt1*-overexpressed RAW264.7 cells with GANT58 and then induced these cells by RANKL. After RANKL induction, we found that the number of osteoclasts among *Dnmt3a*^OE^ cells was more increased than that in WT cells (**Fig. 6h**), while in *Dnmt1*^OE^ cells, osteoclast formation was still inhibited (**Fig. 6i**). We analyzed the osteoclast formation- and function-related proteins NFATc1, CTSK, and MMP9 and found that GANT58 reduced the expression of these proteins. However, these proteins were upregulated when DNMT3a was overexpressed (**Fig. 6j-m**). These findings suggested that DNMT3a might be downstream of GLI1 during the regulation of osteoclast formation.

**Figure 6.**
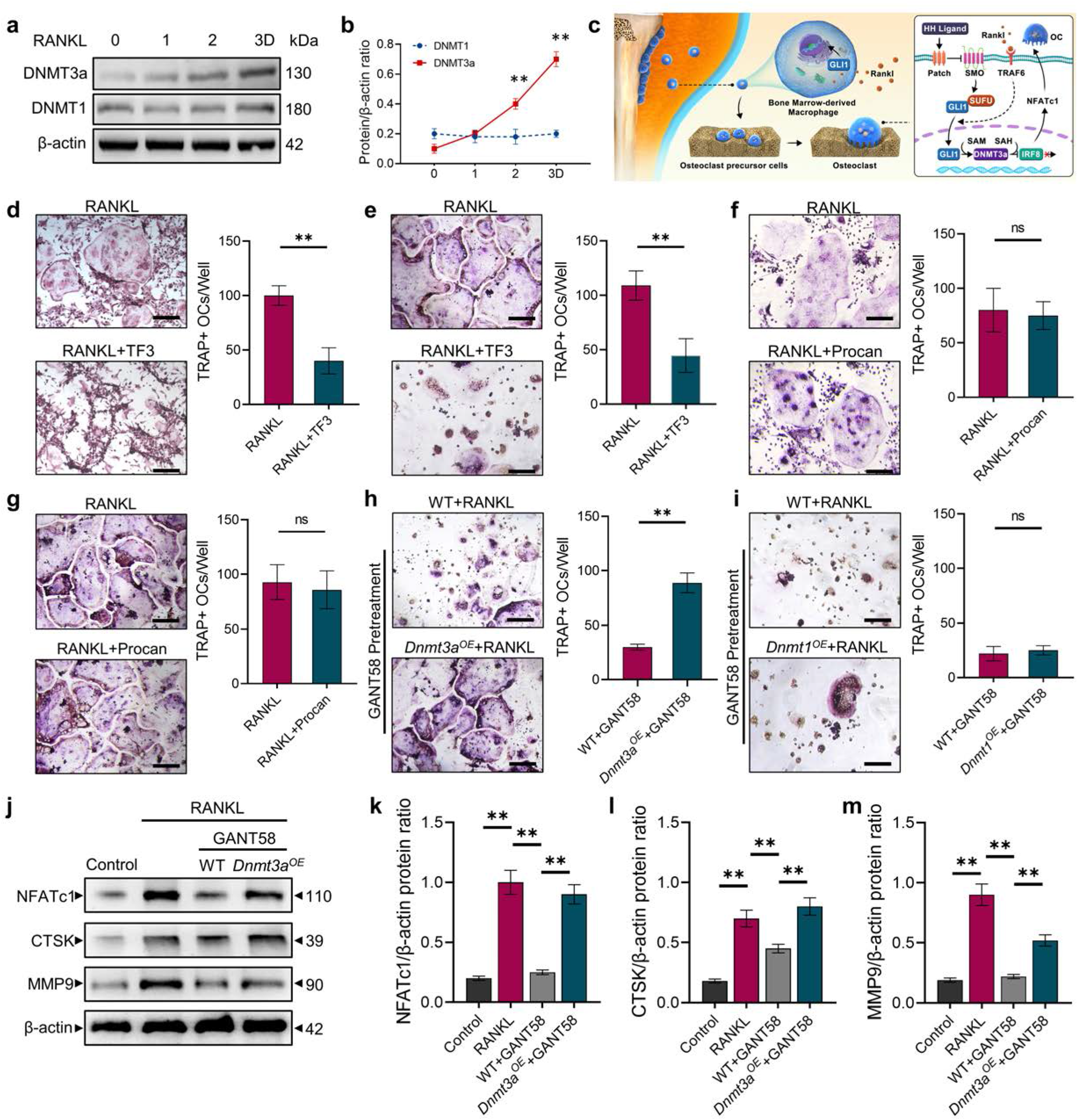
GLI1 regulates DNMT3a to affect the differentiation of osteoclasts. **a** RAW264.7 were stimulated by RANKL for 3 days. Western blot analysis of DNMT3a and DNMT1 during the process of osteoclast induction and **(b)** greyscale value ratio to β-actin of western blot results. n = 3. **c** Schematic diagram of the regulation mechanism. **d, e** TRAP staining of BMMs and RAW264.7 cells stimulated by RANKL (50 ng/ml) in the presence or absence of TF3 (2 μM) and TRAP-positive osteoclast quantity per well. Scale bars = 25 μm. **f, g** TRAP staining of BMMs and RAW264.7 cells stimulated by RANKL (50 ng/ml) in the presence or absence of procainamide (10 μM) and TRAP-positive osteoclast quantity per well. Scale bars = 25 μm. **h** Wild-type and Dnmt3a overexpression RAW264.7 cells were stimulated by RANKL in the presence of GANT58 intervention. TRAP staining and TRAP-positive cell quantity per well. Scale bars = 25 μm. **i** Wild-type and Dnmt1 overexpression RAW264.7 cells were stimulated by RANKL in the presence of GANT58 intervention. TRAP staining and TRAP-positive cell quantity per well. Scale bars = 25 μm. Statistical analysis was performed using Student’s t test. **j** Western blot results of NFATc1, CTSK and MMP9 protein expression. **k-m** Greyscale value ratio to β-actin of western blot results. n = 3. Statistical analysis was performed using one-way ANOVA test. Data shown represent the mean ± SD. *p < 0.05, **p < 0.01, ns = not significant.

## 3. Discussion

In the development of RA, synovitis infiltration and osteoclast overactivation are the direct causes of joint damage [1]. To find a better treatment strategy, we hope to identify a key target that can regulate inflammatory bone destruction using its specific inhibitor and clarify its specific mechanism to carry out targeted interventions and achieve the desired therapeutic effect. Macrophages are an important source of inflammatory cytokines and the primary cells associated with osteoclast formation [17]. In RA, M1 macrophage polarization is generally considered to be an important factor that promotes the development of inflammation [18]. The release of the proinflammatory cytokines IL-1β, IL-6 and TNF-α promotes the activation of more osteoclasts and leads to the destruction of bone structure.

GLI1 is a downstream transcription factor of the hedgehog-GLI signaling pathway. After hedgehog is activated, GLI1 and its factors form a complex with microtubules, enter the nucleus and activate downstream gene transcription [13]. In previous studies, GLI1 signal transduction and other pathways, including the NF-κB signaling pathway, were usually studied in tumor-associated diseases and are considered a response network that promotes cancer development [19, 20]. However, the research results of the hedgehog pathway in bone metabolism are complex. In Heller’s report, enhanced hedgehog signaling due to PTCH heterozygosity indirectly enhanced tumor-induced bone resorption [21]; Ling also demonstrated that the inhibition of GLI1 maintained bone mass [22]. These results showed that GLI1 had a potential connection with the formation of osteoclasts. In contrast, Shoko Onodera concluded that impaired osteoblastogenesis was restored to normal levels by treatment with a small molecule that actives hedgehog signaling [23]. This diversity may be due to the unknown complex mechanism of various factors in the activation of the hedgehog signaling pathway. In our study, we found that GANT58 had an obvious inhibitory effect on the osteoclastic induction of BMMs and RAW264.7 cells, which was consistent with the results of Heller’s study. This seems to mean that the activation of GLI1 may have multiple effects on the regulation of bone tissue homeostasis and maintain a balance in the physiological state. Inflammation is an important reason for osteoclast activation [24]. The release of proinflammatory cytokines promotes arthritis and osteoclastogenesis [25, 26]. Our study showed that GANT58 inhibited the formation of M1 macrophages in an inflammatory environment and inhibited the expression of the proinflammatory cytokines IL-1β, IL-6 and TNF-α. Interestingly, in previous studies, it was found that GLI1 did not appear to respond very strongly to LPS intervention alone [27-29]. In this study, GLI1 played a positive role in the formation of M1 macrophages, and at this point, the role of IFN-γ may be more important. Studies have reported that simple IFN-γ stimulation promotes the expression of GLI1 in neuron precursor cells [30, 31]. These results seem to further confirm that the immunomodulatory role of GLI1 in RA is mainly achieved by macrophage polarization changes. In subsequent studies, we may be able to further explore the regulatory effects of LPS and IFN-γ on GLI1, respectively.

To clarify the downstream regulatory mechanism of GLI1, bioinformatics prediction analysis and RNA-seq were performed and ultimately identified the downstream DNMT family. In addition to normal physiological development, the abnormal expression of DNMTs causes the development of tumors and other diseases [32]. Previous reports have suggested that the absence of DNMT3a inhibits the formation of osteoclasts, which may be due to the methylation of downstream IRF8 by DNMT3a [33]. In our study, we also verified this finding with TF3 intervention. Similarly, in myeloma disease models, high expression of DNMT3a is thought to promote hypermethylation of Runx2, osterix and IRF8 CpG islands and inhibit the expression of these genes, thus inhibiting osteoblasts and promoting osteoclast activation [34]. In contrast to DNMT3a, the overexpression of *Dnmt1* promotes proinflammatory cytokine production in macrophages and plasma during atherosclerosis and inflammation [35, 36]. Notably, GANT58 significantly inhibited DNMT1 and DNMT3a activation during M1 macrophage and osteoclast induction, overexpression of *Dnmt3a* and *Dnmt1* reversed the inhibitory effect of GANT58 on the formation of osteoclasts and M1 macrophages. According to the sequencing and research results, GLI1 activity is closely to affecting the expression activity of DNMTs. In the cell resting state, we found that the inhibitory effect of GANT58 on DNMTs was limited to DNMT3a at the level of gene transcription, while Co-IP analysis found a binding relationship between GLI1 and DNMT1, but not DNMT3a. This evidence suggested that the active regulation of GLI1 on DNMT1 might be post-protein translation, while the regulation of DNMT3a’s expression might be located at the gene transcriptional level. The molecular function of GLI1 usually acts as a transcriptional activator. Patricia Gonz á lez Rodr í Guez’s latest research shows that during autophagy induction, a selective GLI1 enrichment can be observed in the regions closer to the Transcription Start Site (TSS) of the DNMT3a gene [37]. This founding supports our speculation that GLI1 may play a regulatory role by combining with the DNMT3a promoter region sequence.

In vivo experiments using the GLI1 inhibitor GANT58 showed its anti-inflammatory role and protection of bone mass. GANT58 showed good therapeutic effects on CIA mice and inhibited the number of osteoclasts around bone tissue. These results not only elucidated, to a certain extent, the molecular mechanism of GLI1 influencing the pathological process of RA, but also showed the potential therapeutic effect of GANT58 on inflammatory bone destruction in RA, which might have clinically translatable research implications. Although we have demonstrated that the inhibition of GLI1 by GANT58 can reduce the inflammatory response and inhibit osteoclast formation and that this mechanism is achieved through the downregulation of DNMTs, these findings also raise new questions. Inflammatory bone remodeling in RA is a complex process. In addition to macrophages and osteoclasts, the functions of synovial fibroblasts and osteoblasts play essential roles in the RA microenvironment. These cells are also closely linked to each other. Synovial fibroblasts OPG and RANKL secreted by osteoblasts are important factors that regulate osteoclasts. Therefore, in a follow-up study, we will extend the study of GLI1 to its regulatory mechanism in osteoblasts.

## 4. Materials and Methods

### 4.1. Experimental animals and human synovial tissue

Male DBA mice aged 6-8 weeks and weighing 15-20 g were randomly selected and fed in a specific pathogen-free (SPF) environment at a room temperature of 25°C, a relative humidity of 60%, and 12 hours of alternating light. All animal experiments were approved by the Animal Ethics Committee of the Soochow University (201910A354). The animals were divided randomly into groups (6 per group): sham group, vehicle control group (CIA model mice treated with phosphate-buffered saline, PBS), and GANT58 (GLI1 specific inhibitor; MedChemExpress, New Jersey, USA) group (mice treated with 20 mg/kg GANT58) or 5-AzaC (DNMTs specific inhibitor; MedChemExpress) group (mice treated with 2 mg/kg 5-AzaC). An emulsion of bovine type II collagen (Chondrex, Redmond, WA, USA) and an equal amount (1:1, v/v) of complete Freund’s adjuvant (Chondrex) was prepared to establish the CIA mouse model. First, 0.1 ml of the emulsion was injected intradermally into the base of the tail on day 0. On day 22, 0.1 mg of bovine type II collagen mixed with incomplete Freund’s adjuvant (Chondrex) was injected. On day 49, all mice were sacrificed (in accordance with the guidelines of the Animal Welfare and Ethics Committee of the Soochow University) for the collection of specimens. Normal human synovial tissues were donated from surgical patients who have non-inflammatory knee joint diseases such as injury in traffic accidents or traumatic fractures. RA synovial tissues were obtained from RA patients. All included patients were scored DAS28 as described [38]. The samples were obtained with the informed consent of the patients, and all operations were approved by the Ethics Committee of the First Affiliated Hospital of Soochow University ((2018) Ethical Approval No.012).

### 4.2. Immunohistochemical (IHC) staining

Tissue samples were dewaxed and rehydrated gradually with xylene and gradient alcohol, and then citrate buffer solution was added for antigen repair. After the samples were washed with PBS, serum was added to the tissue surface and incubated for 30 minutes at room temperature. The required primary antibody dilutions (GLI1, 1:200; DNMT1, 1:200; DNMT3a, 1:200; Abcam, Cambridge, UK) were prepared, added to the samples after the serum was removed, and incubated overnight at 4 °C. After the samples were washed with PBS, biotin-labeled secondary and antibody working solutions (VECTOR, Burlingame, CA, USA) were gradually added to the tissue samples and incubated for 30 minutes separately. A controlled diaminobenzidine (DAB; Cell Signaling Technology, Danvers, MA, USA) color reaction was performed under a microscope, and the reaction was stopped with distilled water immediately after color development. Hematoxylin staining was then performed. Finally, the samples were immersed in an 80% ethanol solution, a 95% ethanol solution and anhydrous ethanol successively for dehydration. Images were visualized optical microscopy and positive stained cells were assessed by Image J software (Version 1.8.0.112).

### 4.3. Micro-CT analysis

The fixed bone samples of mice were collected. The joint samples were placed in a SkyScan 1174 Micro-CT scanning warehouse (Belgium). The parameters were set as follows: voltage 50 kV, current 800 μA, scanning range 2 cm × 2 cm, and scanning layer thickness 8 μm. The scan data were then entered into the computer to conduct three-dimensional reconstruction with NRecon software, and the bone tissue parameters were analysed with CTAn software after reconstruction.

### 4.4. Hematoxylin and eosin (H&E) staining

The fixed murine bone tissues were removed, placed in 10% ethylenediaminetetraacetic acid (EDTA, Sigma, St. Louis, Missouri, USA) for 3 weeks and paraffin-embedded and sectioned. Other tissues were paraffin embedded and sectioned after being fixed directly. The tissue sections were placed into xylene to dissolve the wax. The sections were then rehydrated in an ethanol solution. After being washed, the sections were immersed in hematoxylin dye (Leagene, Beijing, China) for 3 minutes. Color separation with 1% hydrochloric acid in ethanol and ammonia was then performed. The sections were then immersed in eosin dye (Leagene) and dehydrated with ethanol and xylene after being washed. Bone erosion and fibrosis were visualized and assessed by optical microscopy.

### 4.5. Cell culture

Murine BMMs and RAW264.7 cells were used in this experiment. BMMs were extracted from the long bones of the lower limbs and cultured in α-minimum essential medium (MEM; HyClone, California, USA) containing 10% fetal bovine serum (FBS; Gibco, California, USA), 30 ng/ml macrophage colony stimulating factor (M-CSF; R&D Systems, Minnesota, USA) and 100 U/ml penicillin/streptomycin (P/S, NCM Biotech, Suzhou, China). RAW264.7 cells were cultured in DMEM (HyClone). All cells were cultured in a 37 °C standard environment and stored at -80 °C with serum-free cell freezing medium (NCM Biotech, Suzhou, China).

### 4.6. Cell viability

Cell viability was assessed by a cell counting kit 8 (CCK-8; ApexBio, Houston, USA) assay according to the manufacturer’s protocol. The inhibition rate was calculated as follows: inhibition rate (x) = (OD_control_ - OD_x_)/OD_control._

### 4.7. Tartrate-resistant acid phosphatase (TRAP) staining

A TRAP staining kit (Sigma) was used in this experiment. The dye solution was prepared according to the instructions. For tissue staining, after the tissue sections were dewaxed, a low concentration ethanol solution was added to rehydrate the sections. The repair solution was added to the surface of the tissue for antigen repair. Then, TRAP solution was added and incubated in the dark for 1 hour. For osteoclast induction, BMMs and RAW264.7 cells were seeded in 24-well plates at densities of 4 × 104/well and 6 × 104/well, respectively. 50 ng/ml of RANKL was added into the medium. For cell staining, the cells were first fixed with 4% paraformaldehyde for 15 minutes and then incubated with TRAP dye solution for 40 minutes. The positive cells were observed under an inverted optical microscopy.

### 4.8. Immunofluorescence staining

BMMs were seeded in 24-well plates at a density of 4 × 104/well with stimulation with RANKL (50 ng/ml) or LPS (100 ng/ml) + IFN-γ (20 ng/ml). After culturing and stimulation, all cells were fixed with 4% paraformaldehyde for 15 minutes. To fully bind the antibody to the antigen, 0.1% Triton X-100 (Beyotime, Shanghai, China) solution was added and incubated on ice for 10 minutes. Then, the cells were washed with PBS 3 times, and blocking buffer was added to the cells for 1 hour. The cell supernatant was removed, and the prepared primary antibody solution was added, followed by incubation at 4 °C for 12 hours. The primary antibody was removed, and the fluorescent secondary antibody solution was added and incubated at room temperature for 1 hour. Finally, DAPI dye (1:20; Yuanye, Shanghai, China) solution was added and incubated for 10 minutes, and then the cells were photographed under a fluorescence microscope (Leica, Wetzlar, Germany). Finally, mean fluorescence intensity were acquired and analyzed by Image J software (Version 1.8.0.112).

### 4.9. ELISA

LPS and IFN-γ-induced cell supernatants were collected. The concentrations of the cytokines IL-1β, IL-6, TNF-α and IL-10 in the cell supernatants were measured by ELISA kits (Multi Sciences, Hangzhou, China). The experiments were performed in accordance with the protocol.

### 4.10. Quantitative real-time polymerase chain reaction (PCR)

Total mRNA was extracted using Beyozol reagent (Beyotime). The concentration and purity of the mRNA were assessed by a Nanodrop spectrophotometer. The isolated mRNA (2 μg) was used for reverse transcription PCR to produce cDNA with an iScript cDNA synthesis kit (Bio– Rad). A total of 2 μL of cDNA product was used for subsequent RT–qPCR analysis using SYBR1 Premix Ex Taq (Takara, Dalian, Japan). All primers used in this study are shown in **Table S1**.

### 4.11. Western blotting

Cells were seeded in 6-well plates at a density of 1 × 106/well with stimulation with RANKL (50 ng/ml) or LPS (100 ng/ml) + IFN-γ (20 ng/ml). First, cells were collected to extract total protein, and the BCA (Beyotime) method was used to adjust the protein concentration. Total protein was mixed with 5× loading buffer (Beyotime) and boiled at 95 °C for 10 minutes. The proteins were separated by SDS polyacrylamide gel electrophoresis (SDS–PAGE; EpiZyme, Shanghai, China) based on their different molecular weights. Electrophoresis was performed using Bio–Rad (California, USA) equipment at 180 V for 40 minutes. Then, the proteins were transferred to a nitrocellulose membrane at 350 mA for 70 minutes using membrane transfer equipment (Bio–Rad). The membrane was removed and placed into western blot blocking buffer for 1 hour at room temperature. The diluted primary antibody was placed on the membrane and incubated at 4 °C for 12 hours, and then the corresponding secondary antibody was added and incubated for 1 hour at room temperature. Finally, a chemiluminescence detection system (Bio–Rad) was used to observe the results.

### 4.12. High-throughput sequencing (RNA-seq)

To further screen for differential genes, we first subjected RAW264.7 cells to a 24-hour adaptive culture, followed by the addition of GANT58 at a final concentration of 10 μM to the GANT58 intervention group and cultured for a total of 24 h. After the cell treatment was completed, cells of the control group and GANT58 treated group were collected respectively, and RNA-seq detection and analysis were entrusted to a professional biological company (Azenta Life Sciences, Suzhou, China).

### 4.13. Co-immunoprecipitation (Co-IP)

After culture under the specified conditions, collected the cells to be tested. Pre-cooled RIPA buffer containing protease inhibitor was added and cracked at 4 °C for 30 minutes. Then centrifuged for 15min at 14000g, and immediately transferred the supernatant to a new centrifuge tube. Prepared protein A agarose and washed the beads twice with PBS, then prepared with PBS to 50% concentration. Added 100 μL protein A agarose beads per 1 ml of total protein (50%), shook at 4 °C for 10 minutes. Centrifuged at 4 °C, 14000g for 15min, transferred the supernatant to a new centrifuge tube, and removed protein A beads. Add a certain volume of antibody to 500 μL in total protein, slowly shook the antigen antibody mixture at 4 °C overnight. Added 100 μL protein A agarose beads to capture antigen antibody complexes, slowly shook the antigen antibody mixture overnight at 4 °C. After centrifugation at 14000rpm for 5 seconds, the agarose bead antigen antibody complex was collected, and the supernatant was removed and washed with pre-cooled RIPA buffer for 3 times. Boiled the sample for 5 minutes, electrophoresis the supernatant, and collected the remaining agarose beads.

### 4.14. Statistical analysis

All data are presented as the mean ± standard deviation (SD). Statistical analysis was performed with an unpaired two-tailed Student’s t test for single comparisons with GraphPad Prism 8 (GraphPad Software, CA, USA). One-way analysis of variance (ANOVA) was used to compare data from more than two groups.

## Acknowledgments

This research was supported by the National Nature Science Foundation of China (82072425, 82072498, 82272567, 82072424), the Natural Science Foundation of Jiangsu Province (BE2020666, BK20220059), the Priority Academic Program Development of Jiangsu Higher Education Institutions (PAPD), the Science and Technology Project of Suzhou (GSWS2020121), the Key Project Supported by the Medical Science and Technology Development Foundation, Jiangsu Province Department of Health (H2019024) and the Special Project of Diagnosis and Treatment for Clinical Diseases of Suzhou (LCZX202003).

## Author Contributions

GG and QG participated in the design, experiments and original draft; YZ participated in the design, experimental operation and technical support; WL, WZ, JB, QW, HT and WW participated in experiments and data analysis; ZW, MG, YX and HY provided the administrative support; BL provided the administrative support and designed the study; DG provided the administrative support, designed and funded the study; All authors approved the final version to be submitted.

## Conflicts of Interest

The authors declare that they have no conflict of interest.

## Data Availability

The authors declare that all data supporting the findings of this study are available within this paper and its Supplementary Information.

